# Inhibition of *Streptococcus pyogenes* biofilm by *Lactiplantibacillus plantarum* and *Lacticaseibacillus rhamnosus*

**DOI:** 10.1101/2024.03.12.584618

**Authors:** Alejandro Gómez-Mejia, Mariano Orlietti, Andrea Tarnutzer, Srikanth Mairpady Shambat, Annelies S. Zinkernagel

## Abstract

The human pathobiont *Streptococcus pyogenes* forms biofilms and causes invasive infections, such as pharyngotonsillitis and necrotizing fasciitis. Bacterial biofilms are more resilient to antibiotic treatment and new therapeutic strategies are needed to control biofilm-associated infections, such as recurrent pharyngotonsillitis. *Lactiplantibacillus plantarum* and *Lacticaseibacillus rhamnosus* are two bacterial commensals used for their probiotic properties. This study aimed to elucidate the anti-biofilm properties of *L. plantarum* and *L. rhamnosus* cell-free supernatants (LPSN and LRSN, respectively) on *S. pyogenes* biofilms grown *in vitro* in supplemented minimal medium. When planktonic or biofilm *S. pyogenes* were exposed to LPSN or LRSN, *S. pyogenes* survival was reduced significantly in a concentration-dependent manner and the effect was more pronounced on preformed biofilms. Enzymatic digestion of LPSN and LRSN suggested that glycolipid compounds might cause the antimicrobial effect. In conclusion, this study indicates that *L. plantarum* and *L. rhamnosus* produce glycolipid bioactive compounds that reduce *S. pyogenes* viability in planktonic and biofilm cultures.

## Introduction

*Streptococcus pyogenes* (aka Group A *Streptococcus* (GAS)) is a human pathobiont that colonizes the throat and skin and may cause severe diseases (1) such as necrotizing fasciitis or streptococcal toxic shock syndrome (STSS) with high mortality rates (2-4). GAS is also the most common bacterial cause of pharyngotonsillitis (5). Despite its full susceptibility to many antibiotics, including penicillin, antibiotic therapy alone is often not sufficient to eradicate GAS. After antibiotic treatment up to 30% of pharyngotonsillitis remain symptomatic with GAS still detectable in cultures. One reason for the treatment failure in pharyngotonsillitis is the development of biofilms (6). Biofilm-forming microorganisms are responsible for a large number of difficult-to-treat infections worldwide (7, 8). Bacteria in biofilms are difficult to eradicate, due to insufficient antibiotic penetration into biofilms and the presence of metabolically inactive bacteria inside biofilms (9, 10). Biofilm formation is recognized as a common strategy for both Gram-positive and Gram-negative bacteria to persist and survive host defenses and antimicrobials. Biofilms are characterized by sessile aggregates encased in a self-produced matrix of extracellular polymeric substances and are found on biological and non-biological surfaces (11). It has been suggested that 99% of the world’s bacteria exist in a biofilm state and that biofilm bacteria vary drastically in their physiology, growth rate and gene expression as compared to their planktonic counterparts (12). At least two-thirds of bacterial infections are estimated to be biofilm-related (13). Biofilms provide an enhanced structural defense against biological, physical and chemical stressors (14) and thus biofilm-associated bacteria are difficult to eradicate. As antibiotics often are not sufficient to control biofilm-associated infections, mechanical removal by surgery may be required but is not always possible. Additional strategies to prevent and treat biofilms, such as the use of commensal bacteria aiming to outcompete and inhibit the growth of human pathogens (15-18), recently gained relevance.

Some of the most commonly used probiotics are members of the former genus *Lactobacillus*, recently redistributed in 23 new genera (19). They belong to the normal mucosal microbiota of humans and animals and have rarely been associated with disease. The former *Lactobacillus* spp. exhibit remarkable antimicrobial activity against pathogenic bacteria thanks to the production of bacteriocins, reactive oxygen species, biosurfactants and exopolysaccharides with anti-biofilm activity (20-22). *Lactiplantibacillus plantarum* (LP) and *Lacticaseibacillus rhamnosus* (LR) belong to the human microbiota (23, 24) and are present in food as probiotics. Previous studies have demonstrated their anti-pathogen properties against *Streptococcus mutans, Candida* spp., *Pseudomonas aeruginosa, Staphylococcus aureus, Salmonella* sp., and uropathogenic *Escherichia coli* (20, 25-27). Since recurring GAS pharyngotonsillitis is such a large problem and the probiotics LP and LR are readily available and would represent an alternative for reducing prolonged antibiotic prescription, this study aimed to provide insights into the antibacterial mechanisms of *L. plantarum* and *L. rhamnosus* cell-free supernatants on *S. pyogenes* biofilms.

## Results

### *S. pyogenes* grows and forms biofilms in defined cell culture media supplemented with bacterial medium

In order to mimic physiological nutrient conditions as found in biofilms in the host, GAS was grown in the defined cell culture medium RPMI1640 (RPMI) with and without bacterial medium (Todd-Hewitt Yeast, THY) supplementation. GAS grew in RPMI if supplemented with 1% THY, however a minimum of 3% THY was needed to maintain cell viability beyond 72 hours (**Figure 1A and 1B**). Furthermore, 3% THY was also the minimum supplement necessary for stable bacterial survival for 72 hours *in vitro* in biofilms without renewal of the growth medium (batch condition) (**Figure 1C**). Biofilm formation was observed after three hours of growth as confirmed by the recovery of bacteria after thorough washing followed by sonication (**Figure 1C and 1D**). Additionally, exchanging the growth medium with fresh medium every 24 hours significantly increased GAS biofilm survival at the 72 hours’ time point (**Figure 1D**) (viable GAS counts above 10^6^ CFU/mL) as compared to batch condition (viable GAS counts below 10^6^ CFU/ml). The formation of GAS biofilm in RPMI supplemented with 3% THY was further confirmed by crystal violet staining (**Figure S1**).

**Figure 1.**
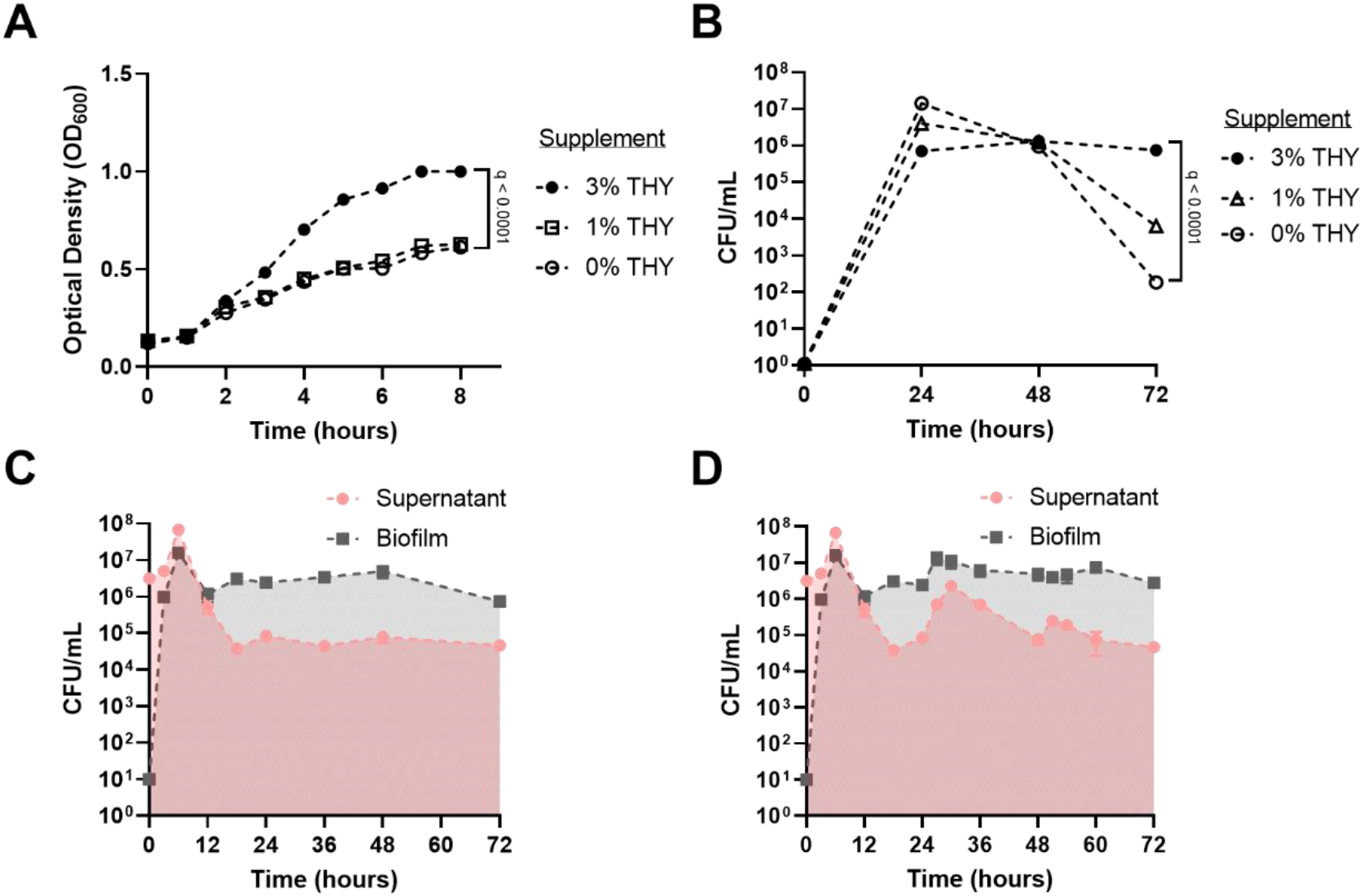
GAS growth and biofilm formation in RPM*I* supplemented with defined concentrations of THY. **(A)** Planktonic GAS growth assessed by optical density (OD_600_) in liquid RPMI supplemented with defined concentrations of THY. **(B)** Planktonic GAS survival in RPMI supplemented with different concentrations of THY. **(C)** GAS biofilm formation and viability in *RPMI* supplemented with 3% THY without medium exchange, determined by CFU/ml. **(D)** GAS biofilm formation and viability in RPMI supplemented with 3% THY and medium exchange every 24 hours, determined by CFU/ml. Data shown as mean ± SD of three independent replicates (n=3). The statistic test used were a non-linear regression model followed by a one-way ANOVA analysis of the growth rate (A) or the maximum population o with Benjamini, Krieger and Yekutieli multiple comparison correction for false discovery rate (**A** and **B**).

### Antibacterial activity of *L. plantarum* and *L. rhamnosus* is medium-dependent

We next assessed the effect of rich bacterial media (THY and MRS) or physiological media (RPMI) on the anti-bacterial capacity of both lactobacilli towards GAS biofilms. Lactobacilli cell-free supernatants prepared using MRS media significantly reduced the viability of a 72-hour-old GAS biofilms (approx. 2.5-log reduction) (**Figure 2A**). In contrast, we observed that cell-free supernatants prepared using RPMI or THY, did not affect the viability of similar GAS biofilms under the same treatment conditions (**Figure 2A**). Next, we determined the effect on the production of lactic acid in the different media as the role of lactic acid, a metabolite produced by *Lactobacillus* spp. is known for its antibacterial properties. The L-lactate content was assessed in all supernatants with the highest content found in MRS supernatants and the lowest in RPMI supernatants (**Figure 2B**). It is also important to note that the content of lactic acid from growth in MRS was similar for both strains of lactobacilli but differed when grown in RPMI or THY, with the highest concentration of lactic acid found in LP supernatants (**Figure 2B**). Adding lactic acid to GAS cell-free spent medium adjusted to pH 4.5 (GASSN-LA) to a concentration comparable to the one present in LPSN and LRSN in MRS (140mM) showed a significant reduction of viable GAS recovered from 72 hour-old biofilms, albeit lower than with MRS derived LPSN or LRSN (**Figure 2C**).

**Figure 2.**
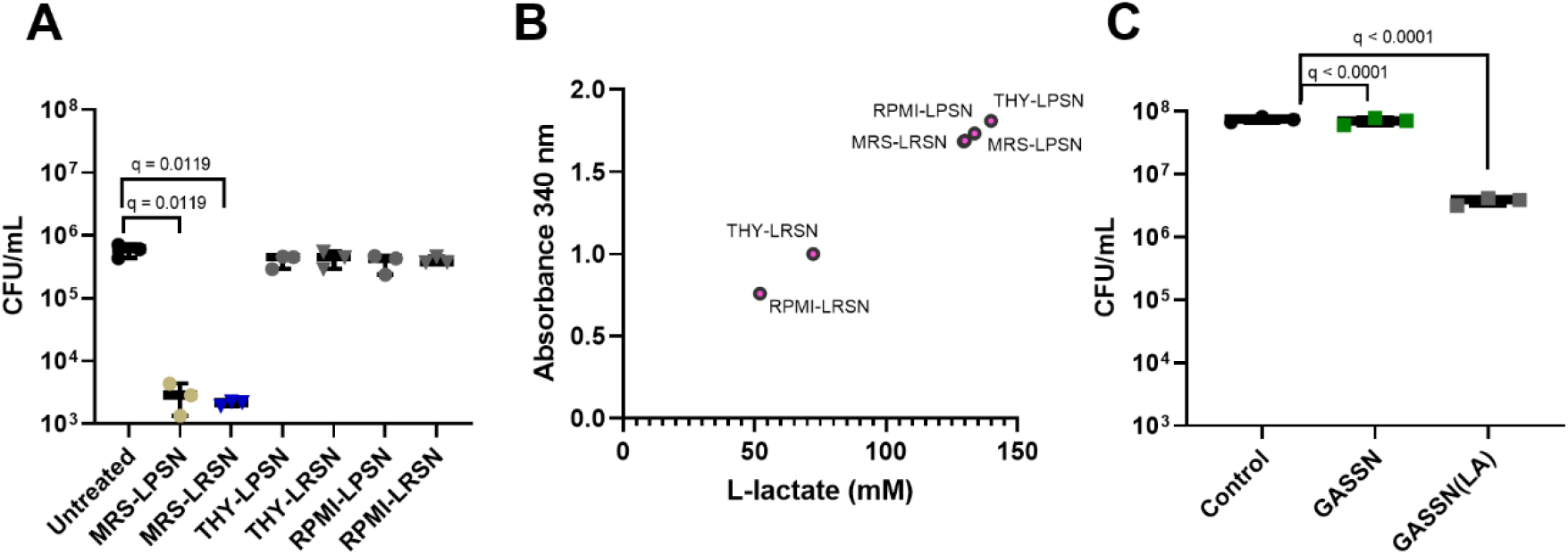
Lactic acid concentration and biofilm treatment with 20% LPSN or LRSN grown in different growth media. (**A**) GAS viable colonies recovered from 72-hour-old biofilms (media exchanged every 24 hours) after treatment with 5 % LPSN or LRSN grown in MRS, THY or RPMI for 48 hours (**B)** fitted values of experimentally determined concentration of L-lactate (mM) from LPSN and LRSN after growth in RPMI, THY and MRS for 24 hours *(red circles)*. (**C**) GAS viable colonies recovered from 72-hour-old biofilms (media exchanged every 24 hours) after treatment with GASSN adjusted with 140mM lactic acid for 48 hours. “control” = untreated, “GA*S*S*N*(LA)” = biofilms treated with GASSN with addition of 140mM lactic acid. For **A** and **C**, biofilms were allowed to form for 24 hours prior to the treatment. Data shown as mean ± SD of three independent replicates (n=3). Statistical significance is indicated as q-values. The statistic test used were an ordinary one-way ANOVA with Benjamini, Krieger and Yekutieli multiple comparison correction for false discovery rate (**A**) and a Kruskal-Wallis with Benjamini, Krieger and Yekutieli multiple comparison correction for false discovery rate (**B**).

### *L. plantarum* and *L. rhamnosus* cell-free supernatants reduce GAS survival in liquid cultures and biofilm in a concentration-dependent manner

The presence of 5% *L. plantarum* cell-free MRS supernatant (LPSN) and *L. rhamnosus* cell-free supernatant (LRSN) in supplemented RPMI cultures led to a significant reduction in GAS bacterial growth (**Figure 3A and 3B**). After 24 hours incubation, a significant reduction in GAS viable cells was observed when treated with either LPSN or LRSN compared to the control group treated with GAS cell-free spent medium with pH adjusted to 4.5 (GASSN) or just MRS medium (**Figure 3A and 3B**). In order to evaluate the anti-GAS biofilm activity of LP and LR, different concentrations of MRS LPSN or LRSN (1, 5, 10 or 20% (v/v)) were added to 24 hours old GAS biofilms. After 48 hours of biofilm treatment (72 hours of experiment) without medium exchange, a significant reduction of viable GAS was observed with 10% LPSN or LRSN compared to the GASSN control (**Figure 3C**). However, when the growth medium was exchanged every 24 hours, a treatment with 20% (v/v) LPSN or LRSN was required to achieve a significant reduction of viable cells (**Figure 3D**).

**Figure 3.**
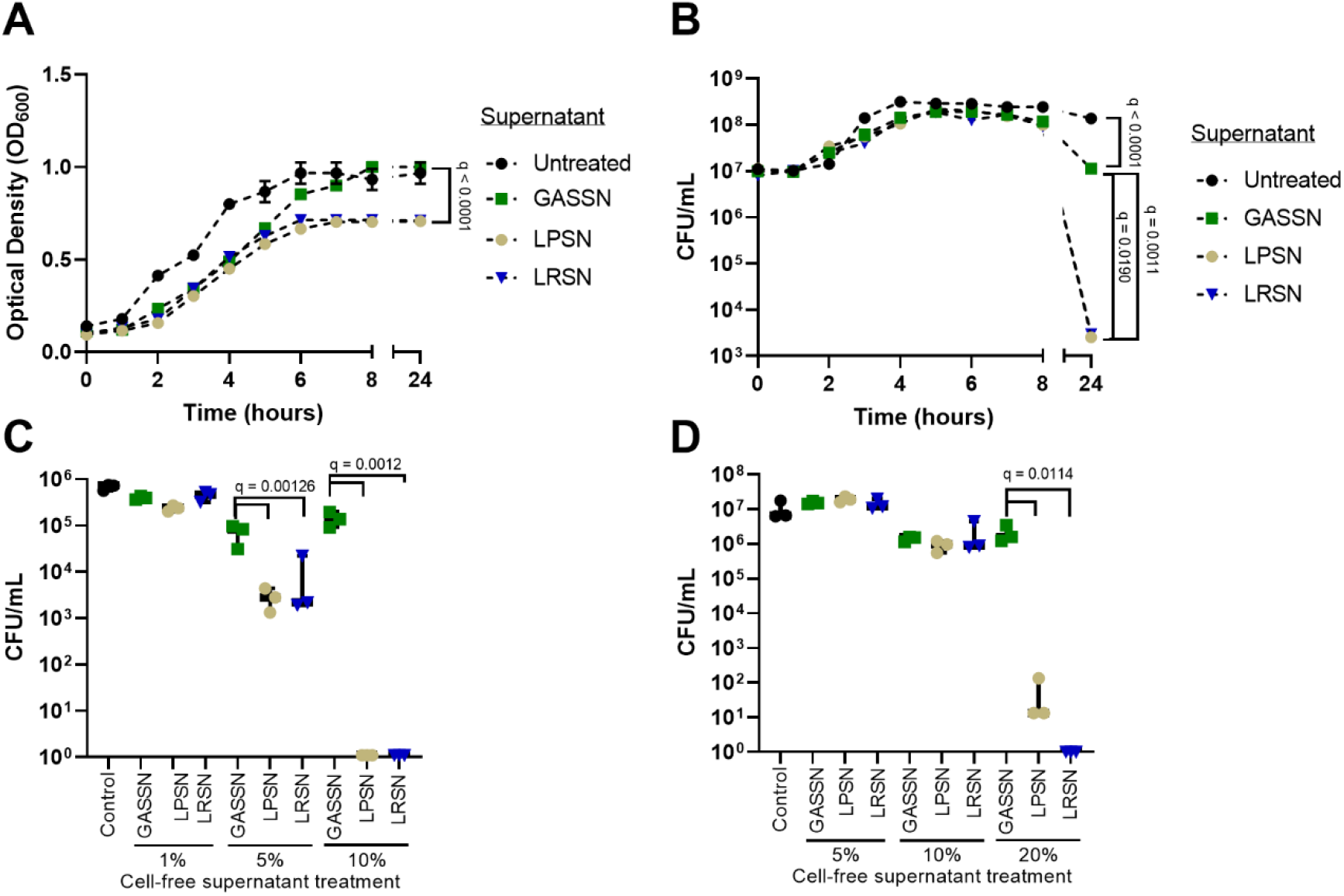
Inhibitory effect of *L. plantarum* (LP) and *L. rhamnosus* (LR) -cell free supernatants on GAS growth in liquid cultures and 72 hours-biofilms. Determination of GAS growth by optical density (OD_600_) (**A**) or as CFU/ml (**B**) after addition of 5% v/v of LPSN (LP) or LRSN (LR) in comparison to GASSN (GAS spent medium adjusted to pH 4.5)). **(C)** Viable GAS CFU/ml counts recovered from 72-hours biofilms without media exchange after 48 hours of treatment. **(D)** Viable GAS CFU/ml counts recovered from 72-hours biofilms with media exchange (every 24 hours) after 48 hours of treatment. For **C** and **D**, biofilms were allowed to form for 24 hours prior to treatment. Control = untreated. Data shown as mean ± SD of three independent replicates (n=3). Statistical significance is indicated as q-values. The statistic test used were a non-linear regression model followed by a one-way ANOVA analysis of the growth rate (**A)** or the population at peak **(B**) and an ordinary one-way ANOVA (**C** and **D**). All cases were analysed with correction for false discovery with Benjamini, Krieger and Yekutieli multiple comparison.

### Efficiency of antibacterial action of LP and LR cell-free supernatants is dependent on the growth state of GAS

Addition of 20% LPSN or LRSN to planktonic GAS cultures before biofilm development (hour 0) did not prevent biofilm formation (**Figure 4A**). However, a decrease in bacterial survival was observed after 24 hours of treatment with an estimate of 48 to 72 hours of treatment required to reduce the number of viable cells to undetectable levels (**Figure 4A**). Corroborating these findings, the LIVE/DEAD BacLight staining imaged by confocal microscope showed a significant increase in damaged or dead cells inside the biofilm reported as the mean intensity of the fluorescence emitted by propidium iodide in both LPSN and LRSN treated groups (**Figure 4B**). Of interest, even though no viable cells were recovered after 72 hours of treatment, a biofilm volume analysis of the confocal images showed a significantly larger biofilm volume after treatment with either 20% of LPSN or LRSN (**Figure 4C**). When evaluating the effect on 24-hour-old biofilms, the addition of 20% LPSN or LRSN led to rapid cell death at 36 hours (12 hours after treatment started) (**Figure 4E**). Microscopic analysis showed a significant increase in the fluorescence signal measured from the staining with propidium iodide after 48 hours of treatment with 20 % LPSN or LRSN (**Figure 4F**). A significant increase in biofilm volume was observed only after treatment with 20% LRSN (**Figure 4G**). 3D figure of scanned biofilms with all colours show the damage caused by the treatments as the intensity of the propidium iodide increases (**Figure 4D and 4H**. 2D representations of all samples can be found in the **Supplementary material (Fig S2 and Fig S3)**.

**Figure 4.**
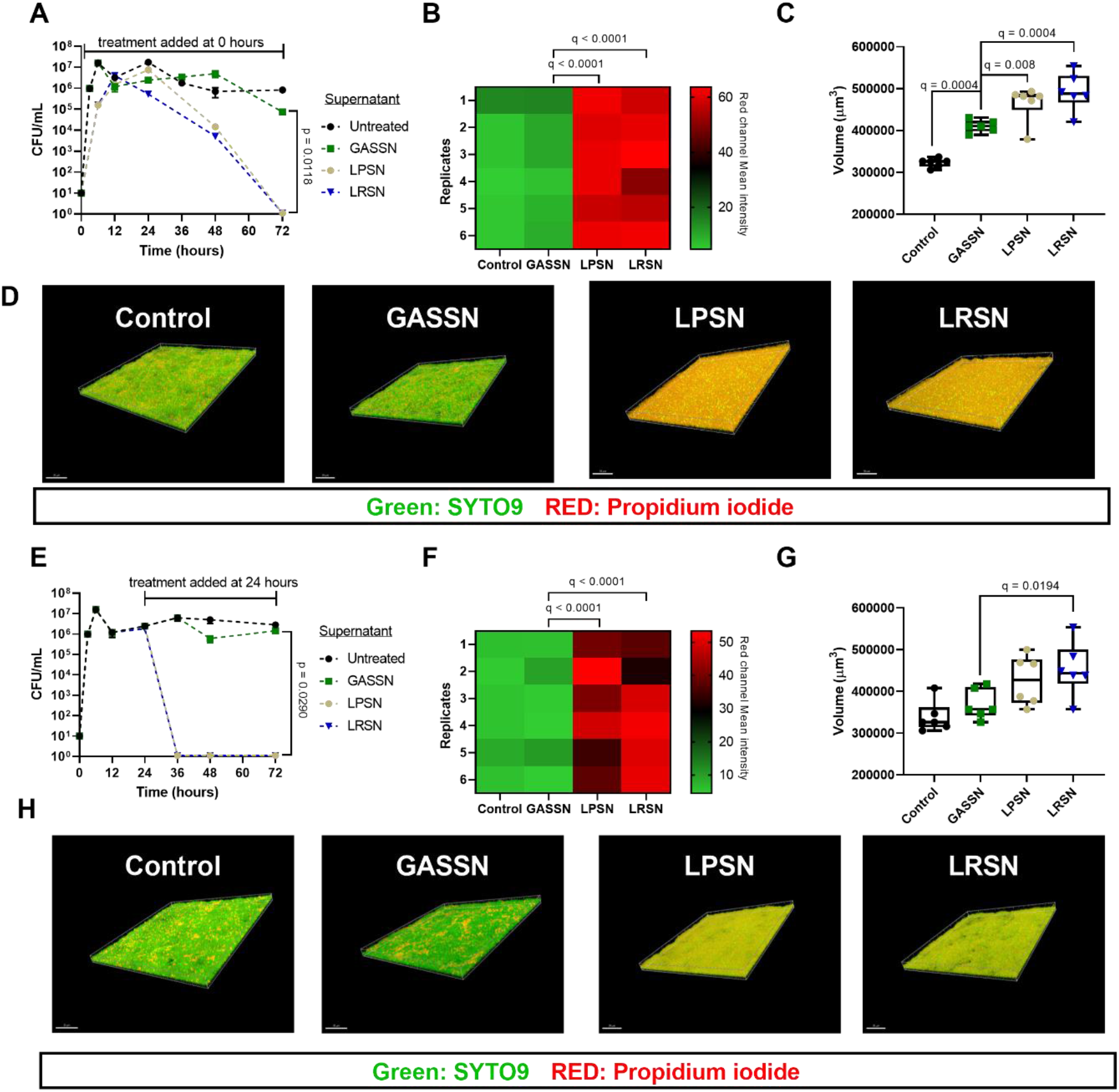
Effect of *L. plantarum* (LP) and *L. rhamnosus* (LR) cell free supernatants on GAS biofilms. **(A-D)** 72 hours of GAS biofilm formation and viability when 0-hours-old biofilms were treated with treated with 20% LPSN or LRSN for 72 hours. **(E-H)** 72 hours of GAS biofilm formation and viability when 24-hours-old biofilms were treated with 20% LPSN or LRSN for 48 hours. **(A and E)** Viable GAS log_10_ CFU/ml counts recovered at 72 hours of experiment. Data shown as mean ± SD of three independent replicates (n=3). **(B and F)** GAS biofilm mean intensity of red (propidium iodide) fluorescence channel after measurement in confocal microscopy at 72 hours of experiment. **(C and G)** GAS biofilm volume in μm^3^ measured with the image scale values at 72 hours of experiment. Data shown as mean ± SD of two representative slide-spots of three independent replicates (n=6). **(D and H)** Representative 3D merged-channels of GAS biofilm models at 72 hours-time point from each treatment group. Scale 30 μm. Statistical significance is indicated as p or q-values. The statistic test used were an ordinary one-way anova (**A and E**) or a mixed-effect analysis (**B, C, F and G**) with Benjamini, Krieger and Yekutieli multiple comparison correction for false discovery rate.

### A glycolipidic compound drives the antibacterial capacity of *L. plantarum* and *L. rhamnosus* cell-free supernatants towards GAS biofilms

In order to categorize potential extracellular metabolites responsible for the antibacterial activity of LPSN and LRSN against GAS biofilms, single and double enzymatic treatments of LPSN and LRSN with proteinase K, α-amylase and lipase were performed. The highest loss of the inhibitory capacity was observed upon lipase treatment for both LPSN and LRSN supernatants (**Figure 5A**). A significant reduction of the inhibitory capacity was also observed upon treatment with a combination of lipase and α-amylase for both supernatants (**Figure 5A**). No significant changes were observed upon treatment with proteinase K (**Figure 5A**). Kinetic experiments confirmed the loss of antibacterial activity of MRS LPSN and LRSN treated with either lipase or amylase (**Figure 5B and 5C**) Control experiments were performed to evaluate the stability of the active molecule in the LPSN and LRSN and to evaluate the effect of enzymatic activity from proteinase-K, α-amylase and lipase on the viability of the biofilm directly. A 1-log decrease was observed in the antibacterial activity of both LPSN and LRSN when exposed to 95°C heat for 3 minutes, a step needed to inactivate the enzymes used for the enzymatic digestions and which indicates that the molecule is thermostable (**Figure S4**).

**Figure 5.**
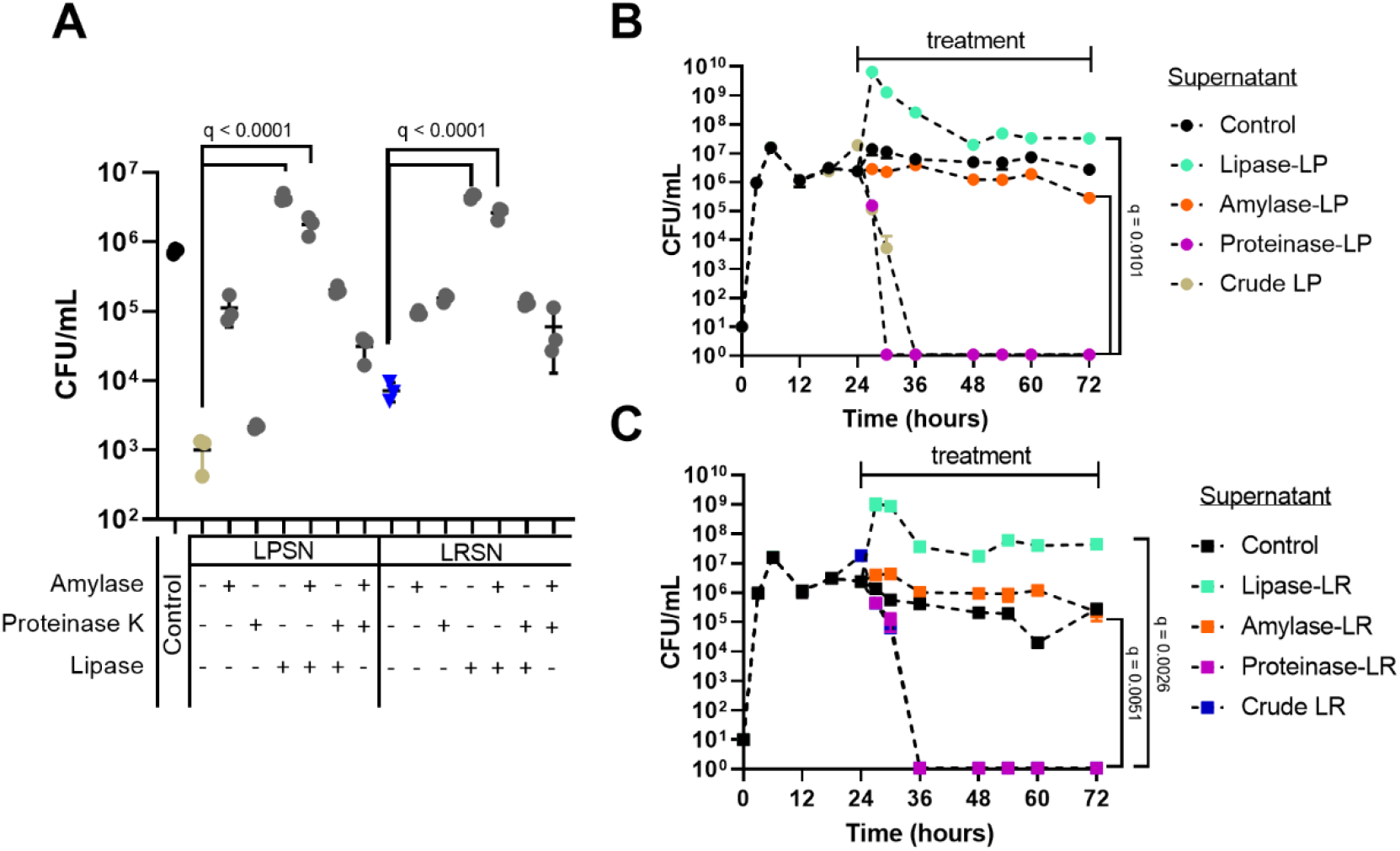
GAS biofilm-inhibitory effect of single- and double-enzymatic treatment of *L. plantarum* (LP) and *L. rhamnosus* (LR) supernatants. 24-hours-old GAS biofilms grown in RPMI supplemented with THY were treated with different digestions (proteinase K, α-amylase or lipase alone or in different combinations) of 20% MRS LPSN or LRSN for 48 hours (with medium exchange every 24 hours). **(A)** Number of viable bacteria recovered from biofilms at the 72 hours time point. **(B-C)** Kinetics of GAS viability in the biofilms under treatment with digested 20% LPSN **(B)** or 20% LRSN **(C)**. GASSN at a concentration of 20% was used as control. Data shown as mean ± SD of three independent replicates (n=3). Statistical significance is indicated as q-values. The statistic test used were an ordinary one-way ANOVA (A) or a mixed-effect analysis (**B and C**) with Benjamini, Krieger and Yekutieli multiple comparison correction for false discovery rate.

Fourier transmission infrared spectroscopy (FTIR) was performed on LPSN and LRSN in order to identify functional groups of organic compounds based on the characteristic infrared absorption peaks of target chemical groups. When LPSN and LRSN grown in MRS were analysed, the main difference was observed at 3200-3600 cm^-1^ spectra, corresponding to OH and ND groups. While in pure MRS the detected signals were weak, a medium OH and a weak NH signal were detected for MRS LPSN and both medium OH and NH signals in MRS LRSN. (**Table 1**).

**Table 1.** FTIR spectrum ranges and functional groups detected in *L. plantarum* and *L. rhamnosus* cell-free supernatants grown in MRS (MRS LPSN, MRS LRSN) compared to fresh growth medium (MRS Control).

| Spectral range ($\text{cm}^{-1}$) | Functional groups detected | | |
| --- | --- | --- | --- |
|  | MRS Control | MRS LPSN | MRS LRSN |
| Below 1000 | CN strong stretching | CN weak stretching | CN weak stretching |
| 1000-1300 | C-O strong stretching; OH strong deformation | C-O strong stretching; OH weak deformation | C-O strong stretching; OH strong deformation |
| 1400-1460 | $\text{CH}_2$ and $\text{CH}_3$ medium asymmetric deformation | $\text{CH}_2$ and $\text{CH}_3$ weak antisymmetric deformation | $\text{CH}_2$ and $\text{CH}_3$ medium deformation |
| 1490-1540 | NH strong deformation | NH strong stretching | NH strong combination |
| 1675-1725 | C=O strong stretching | C=O strong stretching | C=O strong stretching |
| 3200-3600 | OH and NH weak stretching | OH medium; and NH weak stretching | OH and NH medium stretching |

## Discussion

In this study, we showed that GAS formed biofilms *in vitro* in well-defined chemical medium when supplemented with THY. We further showed that the treatment of GAS biofilms with *L. plantarum* and *L. rhamnosus* supernatants reduced the number of viable GAS in the biofilm significantly. This anti-biofilm effect was more pronounced when the cell-free supernatants were added to biofilms as compared to planktonic bacteria. The anti-GAS biofilm property of LP and LR cell free supernatants is possibly bound to a variety of bioactive compounds of the lipids or glycolipids class produced by these two commensal species.

An overall antibiotic failure rate of 20 to 40% has been reported for GAS (28), although GAS antibiotic susceptibility studies have reported no resistance to penicillin so far and the mechanism for this resistance remains elusive (29). In addition, antibiotic resistance to macrolides, tetracycline and fluoroquinolones has been reported (30). Thus due to evolving antibiotic resistance as well as the intrinsic antibiotic tolerance of bacteria in biofilms, new infection control approaches are needed. Biofilm-related infections represent a challenge in terms of infection control. Biofilm-like *S. pyogenes* communities have been observed in tonsillar reticulated crypts, suggesting a role for biofilm in asymptomatic *S. pyogenes* carriage (3, 31).

Cell culture media are well-defined minimal media used to culture eukaryotic cells but not usually bacteria, which are grown in extremely rich and not well-defined media. Previous studies showed that the mucosal pathobionts *Candida albicans, Staphylococcus aureus* and *Pseudomonas aeruginosa* form *in vitro* biofilms in the tissue culture medium RPMI (32-34). Aiming to use a physiological medium that mimics the composition of human fluids (35), we assessed GAS growth, biofilm formation and survival in RPMI. To our knowledge, the *in vitro* growth and biofilm formation capacity of GAS in supplemented RPMI has not been investigated before. Supplementation with THY allowed bacterial growth and survival in liquid cultures as well as biofilm formation for the duration of the experiments performed in this study (72 hours). The decreased biofilm viability in RPMI alone or supplemented with less than 3% THY might be due to lack of nutrients.

The antibacterial effect of probiotic bacteria on important human pathogens such as carbapenem-resistant *Enterobacteriaceae* (36), *Salmonella enterica, Staphylococcus aureus* and *Candida albicans* (37, 38) were shown in previous studies (20). Probiotics include commensal bacteria that display several antibacterial properties, including the former genus *Lactobacillus* (39). For the case of *Lactobacillus* antibacterial activity, a significant effect is attributed to the production of lactic acid as a growth inhibitory substance (40-44). Here we that the loss of both GAS planktonic and biofilm viability was reduced more by treatment with LP or LR supernatants than GAS-spent media adjusted with lactic acid, suggesting that lactic acid was not the main cause for the antibacterial activity. Additionally, we observed that the presence of LPSN or LRSN during the initial stages of biofilm formation did not prevent cell attachment. Nevertheless, the treated biofilms were not viable anymore after 72 hours exposure to LPSN or LRSN. This could indicate that planktonic GAS are less susceptible to the treatment with LPSN and LRSN than those already embedded in a biofilm. Similar findings were described before for other *Streptococcus spp*. (45, 46), suggesting that environmental low pH can enhance biofilm formation as an adaptive response. In addition, when pre-formed biofilms (24 hours of growth) were treated with LPSN or LRSN, a drastic and almost instantaneous decrease in biofilm viability was confirmed by a significant reduction in CFU numbers within 6 to 12 hours after treatment. Hence, by treating GAS in its planktonic state and biofilm state, we observed a higher susceptibility of pre-formed biofilm, which was not evaluated in similar studies conducted previously. Consistently with our findings, a significant antagonistic effect of *L. plantarum* on *S. pyogenes* biofilms in the absence of significant acidification or cell-cell contact has been reported (47). Humphrey et al (45) proposed that the antagonistic effect on *S. pyogenes* might be linked to plantaricin, a peptide produced by *L. plantarum*. We however demonstrated that treatment of *L. plantarum* cell-free supernatant with proteinase K does not affects its antibacterial effect on GAS biofilms. Danilova et al. assessed the effect of proteinase K on *L. plantarum* cell-free supernatants and, consistently with our observations, only non-peptide components were responsible for the antimicrobial activity against *E. coli, P. aeruginosa, S. aureus* and *S. pyogenes* (48). *L. rhamnosus* cell-free supernatants were previously shown to inhibit cell-surface components mediating eukaryotic cell invasion by GAS *in vitro* (49). Furthermore, the effect of *L. rhamnosus* on the adherence of *S. pyogenes* to eukaryotic cells was studied, suggesting that effector molecules released by *L. rhamnosus* and other former *Lactobacillus* strains attenuate the production of virulence factors involved in *S. pyogenes* colonization, nevertheless, no specific secreted metabolites were identified (50). Our data suggest that lipidic or glycolipidic compounds may be involved in the observed antibacterial effect, since only the treatment with lipase inhibited the anti-biofilm properties of both cell-free supernatants. Although no specific lipidic compounds have been directly correlated with the antimicrobial activity against GAS, recent studies (51) have reported the presence of lipidic biosurfactants with predominantly elaidic and palmitic fatty acids in *L. plantarum* cell-free supernatant. Our data suggests that the anti-biofilm potential of both LP and LR is not only dependent on the lactic acid production or acidic pH.

Fourier-transform infrared spectroscopy (FTIR) has been used before to characterize the functional groups associated to biosurfactants present in supernatants derived from different species of the former genus *Lactobacillus*. Based on our results, a more intense band detected at 3200-3600 cm^-1^ suggested the presence of carbohydrates or glycoproteins in all LP and LR supernatants when compared to the medium alone. This may be attributed to lipidic ester bonds present in biosurfactants secreted by *L. plantarum* and *L. rhamnosus*. Furthermore, it was reported that *Lactobacillus delbrueckii* and *Lactobacillus helveticus* produce glycolipidic biosurfactants (52, 53). Other lactic acid bacteria produce glycoproteic biosurfactants containing glucose, mannose, fructose, and rhamnose residues (54, 55). Based on the enzymatic treatment and FTIR analysis of LPSN and LRSN, we suggest that both species produce glycolipidic biosurfactants and that these biosurfactants are responsible for the antibacterial effect on GAS. However, this remains a hypothetical suggestion since the purification of the molecule was not achieved in this study and presents an important limitation for the scope of this study. Furthermore, the antibacterial effect was only observed when GAS biofilms were treated with LPSN or LRSN derived from cultures using MRS medium suggesting that the production of the antibacterial molecule by LP and LN is medium-dependent.

Overall in this study, we showed that cell-free supernatants derived from *L. plantarum* and *L. rhamnosus* MRS liquid cultures exhibited a significant anti-bacterial potential against *S. pyogenes* in planktonic as well as biofilms *in vitro*. Although further studies are required, the anti-bacterial agent is suspected to be a lipidic or glycolipidic compound. Thus, confirming that both *L. plantarum* and *L. rhamnosus* produce and secrete bioactive compounds that could have an important therapeutic value, which could potentially be used in the future to reduce GAS biofilms in conjunction with antibiotics.

## Acknowledgement

The University of Zürich Clinical Research Priority Program BacVivo – Precision medicine for bacterial infections grant supported this work.

We would like to acknowledge the group of Markus Niederberger’s for allowing us to use his FTIR device for this study.

## Methods

### Bacteria, media and growth conditions

Liquid overnight and solid cultures of *S. pyogenes* M1T1 5448 strain (56) were grown aerobically at 37°C in Todd-Hewitt Broth supplemented with 2% yeast extract (THY, BD, France). Liquid and solid cultures of *L. plantarum* and *L. rhamnosus* were grown aerobically at 37° in De Man Rogosa and Sharpe (MRS) medium (Merck, Germany).

### Bacteria reconstitution, isolation, and identification

Lyophilized *L. plantarum* Lp 115 SD-5209 and *L. rhamnosus* GG SD-7017 are marketed by Renew Life (UK) in Florabiotic Everyday Plus® capsules. Maximum recovery diluent (MRD, Oxoid, UK) was used as diluent for bacterial reconstitution. Reconstituted bacteria were diluted and plated on MRS and incubated aerobically at 37°C (57). Individual colonies were acquired randomly and spotted onto MALDI-TOF Biotyper target plates. 1 μL of MALDI matrix (10 mg/mL solution of α-cyano-4-hydroxycinnamic acid (HCCA) in 50% acetonitrile/2.5% trifluoroacetic acid) was added onto each spot and dried. MALDI-TOF/Microflex LT (Bruker Daltonics, USA) was used for automatic measurement and data interpretation. Isolates with logs (scores) ≥ 2 were accepted as correct species identification. Accepted colonies were replated and incubated aerobically at 37° before reconfirmation with MALDI-TOF following the same procedure (58). Whole genome sequencing was performed for both isolated strains to confirm species identity.

### Cell-free supernatant preparation

*L. plantarum* and *L. rhamnosus* were cultured aerobically at 37°C for 24 hours in MRS broth, THY broth or Roswell Park Memorial Institute 1640 (RPMI) (Gibco) supplemented with 3% THY. GAS was grown for 24 hours in THY. Bacterial suspensions were centrifuged (3350 g for 10 minutes) and supernatants were filtered with 0.22 μm filters (TPP, Switzerland). Cell-free supernatants of *L. plantarum* (LPSN) and *L. rhamnosus* (LRSN) were stored at 4°C until further use (40). To obtain GAS cell-free spent medium (GASSN), GAS cell-free spent medium pH was adjusted to the same pH as the MRS cell-free supernatants LPSN and LRSN using 12M HCL, to a final pH of 4.5.

### Determination of lactate concentration

Lactic acid concentrations were assessed in *L. plantarum* and *L. rhamnosus* supernatants grown in either MRS, THY or RPMI supplemented with 3% THY for 24 hours. Cell-free supernatants were collected following the previously described procedure. Lactate was quantified using a lactate assay kit (D-Lactic/L-Lactic acid UV method, r-biopharm, Roche). All samples were measured using fresh growth medium as control.

### GAS growth determination and liquid culture inhibition assay

*S. pyogenes* liquid cultures were prepared by dilution of an overnight culture to OD_600_ 0.1 in RPMI (Gibco) supplemented with 3% THY in the presence or absence of 5% LPSN and LRSN. The LPSN and LRSN were prepared in THY, RPMI or MRS, with MRS supernatants chosen for all the biofilm assays. Overnight cultures were washed once with sterile PBS before dilution. Survival rates were determined by colony forming units (CFU/mL) counts every hour (58). RPMI with 3% THY was used as control for GAS growth.

### Biofilm determination by crystal violet staining

Once the incubation time of the biofilms finish, the biofilms were washed with PBS. The liquid was removed and the biofilm were dried for 1h at 50°C. Next, a solution of 0.1% Crystal Violet was added (1:10 diluted in milliQ H2O) and the biofilms were incubated for 30 minutes at RT after which the biofilms were washed with MilliQ H20. The liquid was removed and the stained biofilm was dissolved in 90% EtOH. An aliquot of the solution was taken into a 96-well plate and the absorbance was measured at 570nm.

### GAS biofilm viability assay

GAS overnight cultures were washed with sterile PBS once before diluting to OD_600_ 0.1 in RPMI (Gibco) supplemented with 3% THY. 2 ml aliquots were transferred to 6-well tissue culture plates (TPP, Switzerland) and incubated aerobically at 37°C. LPSN or LRSN (1, 5, 10 or 20% concentrations) obtained from growth in MRS were added after 24 hours of incubation to the pre-formed GAS biofilms. For biofilms grown with medium exchange every 24 hours, LPSN or LRSN were freshly added along with the medium change. GASSN was used as a control for the effect of spent medium and low pH. After 72 hours incubation, the supernatant of each well was collected, biofilms were washed three times with sterile PBS and attached bacterial cells removed mechanically by sonication and scratching of the bottom of the wells. Supernatants and biofilm suspensions were serially diluted, plated on THY agar and incubated aerobically at 37° for CFUs enumeration. Viable counts were expressed as the CFU per mL.

### Treatment of cell-free supernatants

In order to assess the origin of the bioactive compound from LPSN and LRSN grown in MRS, an enzymatic treatment was performed. α-amylase (150 units/mL, Sigma-Aldrich), proteinase K (1 mg/mL, Sigma-Aldrich) and lipase (25 mg/mL, Sigma-Aldrich) were used. The different enzymes were added individually or in combination to both LPSN and LRSN and incubated at 37°C for 3 hours, followed by heat inactivation at 95°C for 3 minutes. The samples were centrifuged (3350 g for 3 minutes) and the supernatant was stored at 4°C until further use (58). Double enzymatic treatments were performed in two steps, as described above. For all conditions, enzyme-treated GASSN was used as control. Adjustment of lactic acid concentration in cell-free supernatants was performed by dilution of a 2M L-lactate stock (Sigma-Aldrich, Belgium) to 140 mM in GASSN.

### Visualisation of GAS biofilms

GAS overnight cultures were washed with sterile PBS once before diluting to OD_600_ 0.1 in RPMI supplemented with 3% THY. Aliquots were transferred to 8-well μ-slides (Ibidi GmbH, Germany) and incubated aerobically at 37°C. MRS LPSN and LRSN were added (20% v/v) at 0 hours or after 24 hours of GAS incubation. Growth medium containing MRS LPSN or LRSN derived was exchanged every 24 hours. GASSN was used as a control. After 72 hours, biofilms were washed three times with sterile PBS, fixed at 50°C for 50 minuftes and stained with LIVE/DEAD BacLight Kit (Molecular Probes-Invitrogen, USA). Biofilms were visualized by confocal laser scanning microscopy (CLSM-Leica SP8 inverse, Center for Microscopy, University of Zurich). Biofilm volume (um^3^) and mean intensity of red channel (dead or damaged bacteria stained by PI) values of three different biological replicates and two technical replicates for each condition were retrieved using the Imaris Microscopy Image Analysis Software (Oxford Instruments, UK).

### Characterization of *L. plantarum* and *L. rhamnosus* cell-free supernatants by Fourier transmission infrared spectroscopy (FTIR)

Cell-free supernatants of *L. plantarum* and *L. rhamnosus* grown in MRS and in supplemented RPMI were obtained as described previously. Functional groups present in the supernatants were analysed using Fourier transmission infrared spectroscopy (FTIR). FTIR spectra for both cell-free supernatants and fresh medium were recorded in the region of 400-4000 cm^-1^ at a resolution of 4 cm^-1^ on an ALPHA FTIR Routine spectrometer (Bruker) (59). Four independent replicates for each condition were analysed. Functional group data analysis was performed on KnowItAll® System 2021 – Spectroscopy Edition (John Wiley & Sons, Inc.).

## Statistical analysis

Statistical analysis of the data was performed with the software Graphpad Prism (v 9.5.1) and the different tests were performed after normality was assessed using the Shapiro wilk test and QQ plots. Growth curves were analysed using a non-linear regression with either Gompertz growth or Beta growth then decay models followed by one-way Anova. Normally distributed data were analysed by ordinary one-way ANOVA while, for non-parametric data Kruskal-Wallis or a Mixed-model analysis was used. All tests were further evaluated post-hoc for multiple comparisons using Benjamini, Krieger and Yekutieli correction for False Discovery Rate. Significance threshold of *p* < 0.05 or q < 0.05 was considered for all experiments.

## Supplementary material

**Figure S1.**
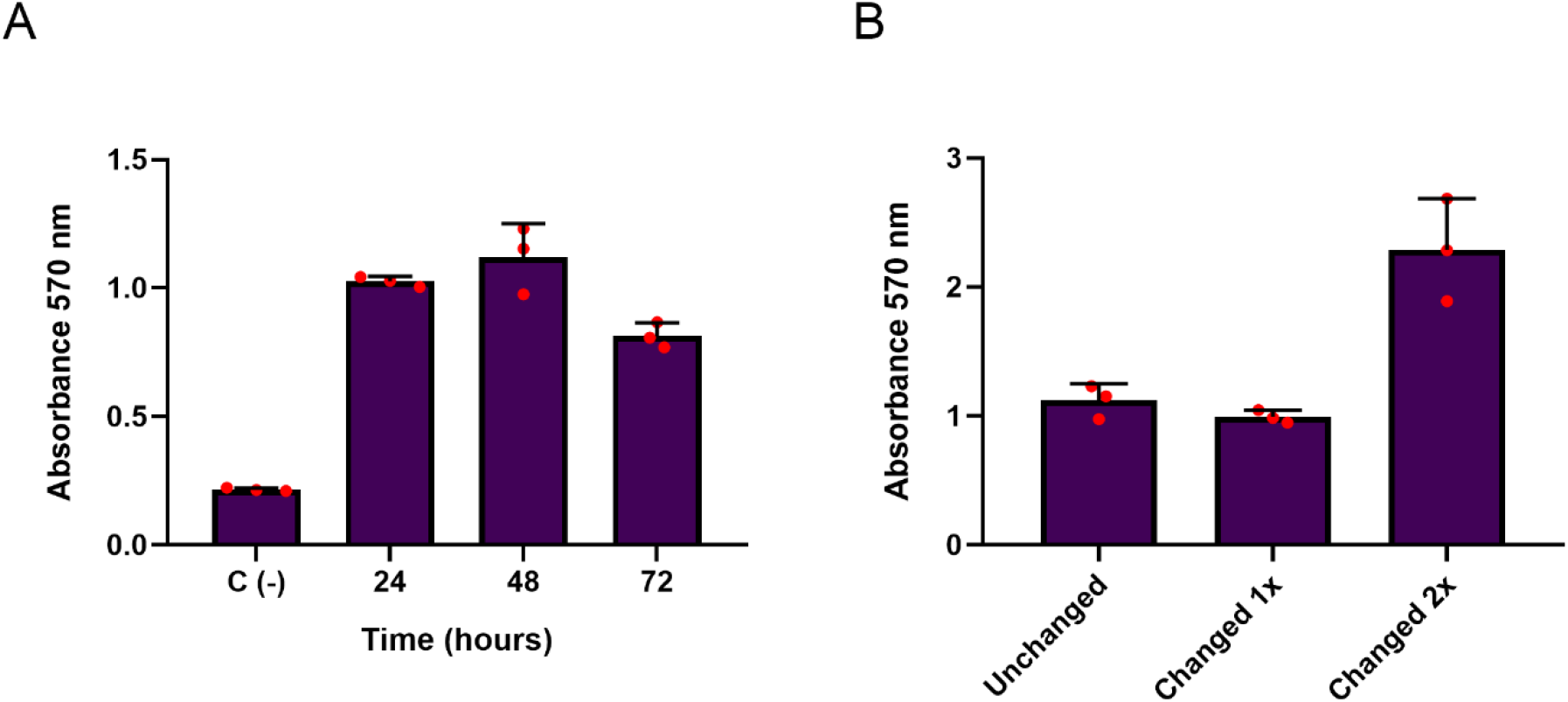
Crystal violet measurements of GAS biofilms in RPMI supplemented with 3% THY. (**A**) OD_570_ measurements of crystal violet staining of GAS biofilms after 24, 48 and 72 hours of static growth without medium exchange. (**B**) OD_570_ measurements of crystal violet staining of GAS biofilms after 72 hours of growth without medium exchange, medium exchange once after 24 hours and medium exchange every 24 hours. Background signal from the crystal violet is shown as C (-). The experiments were performed in biological triplicates.

**Figure S2.**
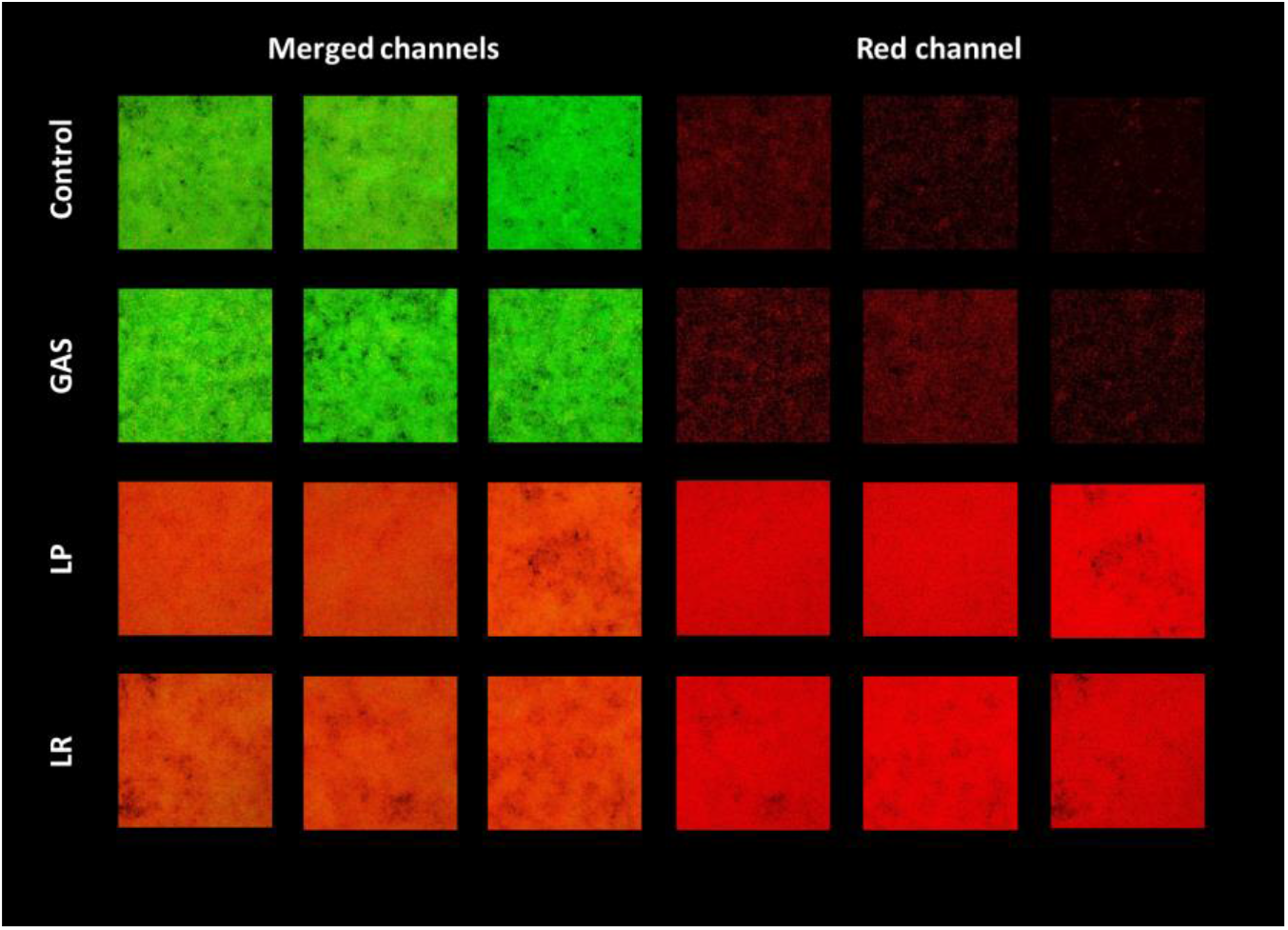
2D models of 72 hours GAS biofilms (n= 3) treated with 20% v/v of MRS LPSN or LRSN at hour 0. “Control” = untreated GAS biofilms. “GAS” = GAS biofilms treated with 20% GASSN. “LP” = GAS biofilms treated with 20% LPSN. “LR” = GAS biofilms treated with 20% LRSN. Biofilms are stained using Syto9 (green channel, live bacteria) and propidium iodide (red channel, dead or membrane damaged bacteria) and imaged with a confocal laser scanning microscope.

**Figure S3.**
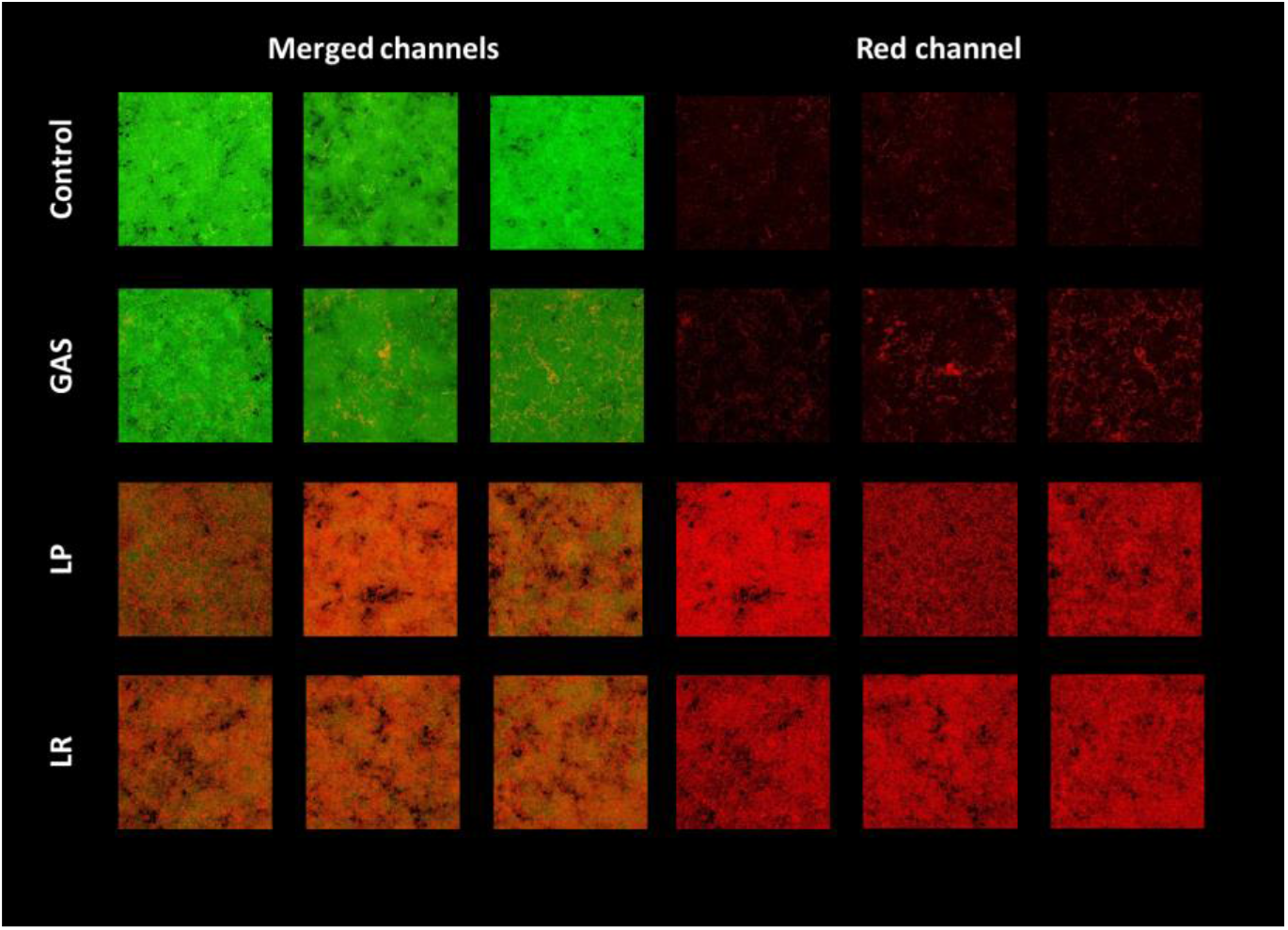
2D models of 72 hours GAS biofilms (n = 3) treated with 20% v/v of MRS LPSN or LRSN after 24 hours of biofilm formation. “Control” = untreated GAS biofilms, “GAS” = GAS biofilms treated with 20% GASSN, “LP” = GAS biofilms treated with 20% LPSN, “LR” = GAS biofilms treated with 20% LRSN. Biofilms are stained using Syto9 (green channel, live bacteria) and propidium iodide (red channel, dead or membrane damaged bacteria) and imaged with a confocal laser scanning microscope.

**Figure S4.**
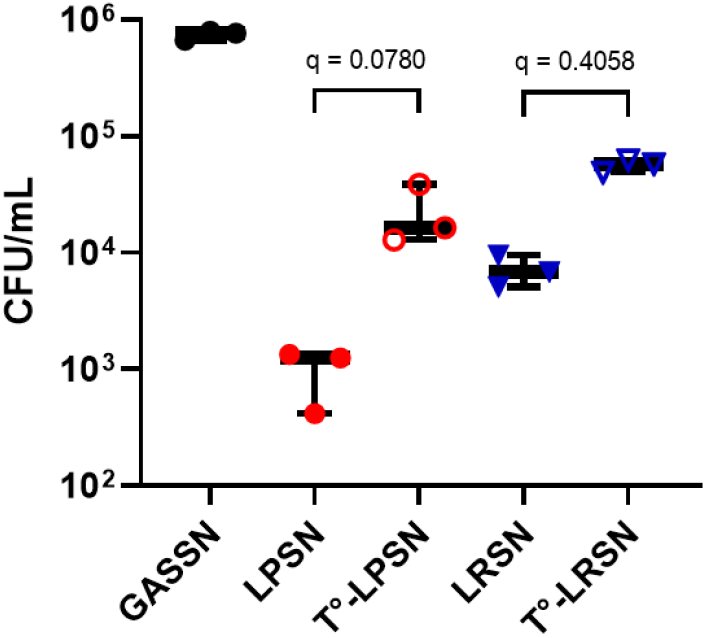
Effect of exposure to 95°C for three minutes on the antibacterial activity of 20% MRS LPSN and LRSN (T°-LP and T°-LR). The prepared supernatants were added to biofilms after 24 hours of growth in culture media supplemented with THY and incubated for a further 48 hours (72 hours total incubation time with media exchange every 24 hours). The CFUs were enumerated for all conditions. Data is shown as mean ± SD of three independent replicates (n=3). Statistical significance is indicated as q-values. The statistic test used was an ordinary one-way ANOVA with Benjamini, Krieger and Yekutieli multiple comparison correction for false discovery rate.

